# Restoring p53 Function and Silencing REV3L Suppresses the Cancerous Metabolic Phenotype in Cisplatin Treated Human Non-Small Lung Carcinoma Cells

**DOI:** 10.1101/200816

**Authors:** Linghao Kong, Michael M. Murata, Michelle A. Digman

## Abstract

Lung cancer is one of the deadliest cancers in the world accounting for over one-quarter of all cancer-related deaths, but in many cases, the cancer can develop a resistance to the cisplatin-based chemotherapies. It is well known that cancer cells exhibit the Warburg effect and some studies have suggested that cancer cell metabolism may be linked to cisplatin resistance. In this study, the effects of tumor suppressor protein p53 and translesional synthesis protein REV3L are studied to relate DNA damage signaling and repair to cellular metabolism by using the fluorescence lifetime of the metabolic coenzyme NADH. It was found that simultaneously restoring function to p53 and silencing REV3L suppressed the cancerous metabolic phenotype and resulted in the greatest amount of cancer cell death. This study demonstrates a previously unrecognized relationship between p53 and REV3L in cellular metabolism and may lead to improvements in chemotherapy treatment plans that reduce cisplatin resistance in cancer cells.

## Introduction

Lung cancer is one of the deadliest cancers in the world. The American Cancer Society estimates that in 2017 alone, over 150,000 people will die of lung cancer, and even though lung cancer only accounts for about 14% of all new cancer cases, it represents 27% of cancer-related deaths [1]. Furthermore, only 3% of patients diagnosed with lung cancer survive the disease making it one of the lowest survival rates for both men and women [2].

Currently, there are several methods for managing and treating lung cancer, including surgery, radiation therapy, and chemotherapy. One of the most common chemotherapy medications used to combat lung cancer is the antineoplastic agent cis-diamminedichloroplatinum (II) more commonly known as cisplatin. In many cases, however, the cancer can develop a resistance to this cisplatin-based chemotherapy. This resistance to cisplatin-based treatment plan endangers the life of the patient, who has an aggressive type of cancer with an ineffective treatment plan.

Cisplatin is normally able to reduce cancer cell proliferation by covalently binding to deoxyribonucleic acid (DNA) and creating intrastrand crosslinks (ICLs) that stall DNA replication forks. The covalent bonds formed in the ICLs are formed most commonly at guanine bases and these sterically inhibit motion of the DNA polymerases at replication forks [3]. The stalling of DNA replication forks occurs when high fidelity DNA replication polymerases cannot add nucleotides across the ICL, causing the polymerases to be ejected. Such stalling is highly toxic to rapidly dividing cells, which is why cisplatin is effective against cancers but has side effects such as hair loss and low blood cell count. The lesions caused by cisplatin activates the DNA damage response (DDR), which in turn activates the tumor suppressor protein p53 which then regulates metabolism, limits proliferation, and resists carcinogenesis [4]. The signaling in the DDR can reduce cyclin-dependent kinase (CDK) activity and this inhibition of CDK can arrest the cell cycle halting cancer cell growth. With continuous exposure to cisplatin, ICLs accumulate in the cell and repeatedly initiate the DDR or prolong signaling. This persistent DNA damage signaling can then induce apoptosis or senescence in the cell, both of which are highly desirable for treating cancers [5].

Several mechanisms of resistance to cisplatin have been proposed, including detoxification by glutathione conjugates, changes in DNA methylation, alterations of membrane protein trafficking, and increased DNA repair [6]. In particular, resistance may be conferred by the DNA repair protein DNA polymerase ζ. In normal DNA polymerase activity, double stranded DNA is separated into individual strands by helicase. DNA polymerases are then able to attach to single strands of DNA and begin assembling nucleotides in the 5’ to 3’ direction. Unlike the highly accurate DNA polymerases used in replication such as DNA polymerase δ, DNA polymerase ζ has low fidelity nucleotide insertion during the translesional synthesis (TLS) DNA repair pathway [7]. Using TLS, cancer cells are able to quickly respond to the lesion site and repair it, thereby overriding DNA damage signaling and continuing proliferation [7]. The reversionless 3-like (REV3L) protein is the catalytic subunit of DNA polymerase ζ that is responsible for TLS [8]. REV3L has been shown to increase cancer cell viability, especially following damage by cisplatin [9, 10, 11].

Another potential mechanism of cisplatin resistance may involve the p53 tumor suppressor protein. DNA damage caused by cisplatin would normally activate p53 to halt the cell cycle or initiate the apoptotic cell death pathway. However, in more than half of all cancers, p53 is either mutated or completely missing [12]. One of p53’s many functions is metabolic regulation, which include the upregulation of oxidative phosphorylation (oxphos) and the downregulation of glycolysis [13]. Without p53, cancers can proceed with unregulated metabolism in the form of high rates of glucose and low rates of mitochondrial oxphos known as the Warburg effect. The Warburg effect describes this preference of glycolysis over oxphos in cancer cells and the current theories as to why this occurs are because glycolysis produces more macromolecules necessary for proliferation and glycolysis can happen without oxygen which is important for cell division in non-vascularized tumors [14]. Interestingly, REV3L may be involved in metabolic regulation, but its effect on metabolism is still debated. Some sources claim that it increases cell reliance on glycolysis and others that it promotes oxphos [10, 15]. Because the hallmark of cancer cells is Warburg metabolism, understanding the mechanisms that limit this cancerous metabolic phenotype may lead to improvements in chemotherapy treatment plans that reduce cisplatin resistance in cancer cells.

In this study, I propose that depleting REV3L and restoring p53 function limits the cancerous metabolic phenotype following cisplatin treatment. H1299 human non-small cell lung carcinoma cells will be used to investigate the metabolic response following manipulation of p53 and REV3L. Metabolism will be tracked using the fluorescence lifetime of an intermediary metabolite to assess relative usage of oxphos and glycolysis. Cell morphology assays will be used to determine whether or not different metabolic phenotypes correspond to decreased viability. The results of this study will help to elucidate the roles of p53 and REV3L in the metabolic regulation of lung cancer cells following a chemotherapy treatment in order to develop more effective treatment plans for cisplatin-resistant cancers.

## Materials and Methods

### Cell Culture and Treatments

This study sought to elucidate the roles of tumor suppressor p53 and the REV3L subunit of DNA polymerase ζ in cellular metabolism following cisplatin treatment of lung cancer cells. Thus, the commercially available H1299 human non-small lung carcinoma cell line (ATCC) was chosen as the system for investigation. In particular, the H1299 cell line has a homozygous partial deletion of p53 and thus, lack expression of endogenous p53 protein. H1299 cells were cultured in RPMI 1640 cell culture media supplemented with 10% fetal bovine serum (FBS), 2 mM L-glutamate, and 1% penicillin-streptomycin. Before imaging, cells were trypsinized and plated with 60-80% confluency on glass-bottom imaging dishes. They were incubated at 37 °C with 5% CO_2_ for 12-24 hours prior to imaging in order to simulate physiologically relevant conditions. During imaging, cells were kept at 37 ^o^C and 5% CO_2_. To induce intrastrand cross-links, cells were treated with 20 μM cis-diamminedichloroplatinum (II) for the duration of the experiments.

### Plasmid Amplification and Transfection

Since H1299 cells do not express functional p53, purified nucleic acid vectors containing the enhanced green fluorescent protein (EGFP), p53 tagged with GFP on the C terminus (p53-GFP), or p53-GFP with the R175H mutation (p53-R175H-GFP) were introduced into the cells via transfection. These nucleic acid vectors contain the p53 protein, an attached fluorescent fluorophore, and an antibiotic region. Because of the attached fluorescent protein, the successful transfection of p53 protein and variants can be visualized using confocal fluorescence microscopy. The EGFP and p53-GFP plasmids are commercially available at Addgene while the p53-R175H-GFP plasmid was kindly constructed and provided by Dr. Lee Bardwell and Dr. Jane Bardwell in the Department of Developmental and Cell Biology at the University of California, Irvine.

To ensure that there was enough plasmid for all experiments, bacterial transformation was used to amplify plasmids. By transforming bacteria with foreign DNA plasmids, the bacterial machinery can replicate the plasmid at a very high rate in a short amount of time thereby saving resources. DH5α competent bacteria, a strain of versatile E. coli, were transformed using the heat shock method, which involves submerging the bacteria in 42 °C water for 1 min and then setting them on ice for 2 min. The sudden heat opens the pores on the bacterial surface to allow plasmids to enter the bacteria. Cooling the bacteria with ice closes the pores quickly so that the bacteria can survive and replicate the plasmid. Then, the bacteria were grown on antibiotic infused agar plates that corresponded to the antibiotic resistance conferred by successful transformation with the desired plasmids. Individual colonies were then selected to ensure that all the bacteria were genetically identical. The colonies were grown in a bacterial tube in an incubator and shaker set to 37 °C and 200 rpm to ensure that the distribution of nutrients is even for all bacteria and to better oxygenate the media at a bacterially preferable temperature. A plasmid purification kit (Qiagen) was used to extract the plasmids through lysing the cells, removing unwanted biological material such as bacterial membranes or ribosomes, and isolating the resulting plasmids using centrifuge-based filtration columns. A Nanodrop 2000c spectrophotometer was used to measure the concentration of isolated plasmids using the Beer-Lambert law, where a greater concentration of plasmid absorbed more light that was passed through the sample.

Transfection of DNA is the process by which a desired protein can be expressed in cells. Through this process, DNA plasmids are encapsulated in lipids, which protects them from degradation and allows for their easy uptake into cells for subsequent transcription and translation. The transfection of DNA plasmids was done using Lipofectamine 3000 delivery system. Imaging was performed 24 to 48 hours after transfection to allow for the expression of plasmid.

By introducing the genetic material for GFP-tagged p53 into the H1299 cells, the effect of functional p53 on cellular metabolism following cisplatin treatment could be determined since H1299 cells do not normally express p53. Transfection with the EGFP plasmid acted as a control for any abnormalities that may arise from the transfection process. Transfection of p53-R175H-GFP served to explore the effect of mutated p53 as commonly found in most cancers in which a point mutation causes a structural change in the p53 protein inhibiting binding to DNA in a sequence specific manner.

### Depletion of REV3L

To evaluate the role of REV3L on cellular metabolism after DNA damage by cisplatin, small interfering RNA for REV3L (siREV3L) or short hairpin RNA for REV3L (shREV3L) were used to silence the gene for REV3L. The siREV3L and shREV3L constructs as well as the empty controls (siControl and shControl) are commercially available at Addgene. The short complementary nucleotide sequences operate within the RNA interference pathway to repress translation of target mRNA sequences by binding to them or marking them for degradation.

The transfections of siREV3L, siControl, shREV3L, or shControl into cells were very similar to the process of DNA transfection. Transfection of siREV3L or siControl was done simultaneously with DNA plasmid transfection and the short interfering RNA was added with the DNA. The process of shRNA transfection occurred after the transfection of the DNA plasmid but the steps were identical, with either shControl or shREV3L used. One day after shRNA transfection, media with 2.0 μg/mL puromycin was applied for 24 hours to select for successfully transfected shRNA cells.

### Tracking Metabolism Using Fluorescence Lifetime of NADH

In this study, cellular metabolism was monitored before and following cisplatin treatment in H1299 lung cancer cells. A key intermediary metabolite in cellular metabolism is the coenzyme nicotinamide adenine dinucleotide (NAD) [16]. NAD is involved in a variety of redox reactions to shuttle electrons between steps in both the glycolytic and oxidative phosphorylation metabolic pathways. Thus, NAD primarily exists in two forms, an oxidized and a reduced form. Reduced NAD (NADH) is particularly interesting because it is auto-fluorescent with 740 nm two-photon excitation, which enables real-time investigation of NADH without the need for labeling. Furthermore, NADH changes its structural conformation from a closed form when freely floating to an open form when bound to a substrate [17]. This conformational change from closed to open causes a change in the fluorescence lifetime decay curve. Fluorescence lifetime is the amount of time between excitation of a fluorophore and emission of fluorescence. Free NADH has a fluorescence lifetime of 0.4 ns while substrate-bound NADH has a lifetime of 3.2 ns. In this way, fluorescence lifetime can be used as a form of contrast that can be used to determine the molecular activity of NADH. Fluorescence lifetime imaging microscopy (FLIM) data was collected using a laser scanning microscope. A Ti:Sapphire Mai-Tai laser is coupled to the imaging system and has a repetition rate of 80 MHz and a pulse duration of approximately 100 fs. Fluorescence images were collected by exciting NADH at 740 nm and collecting the photons with a bandpass filter at 460/80 nm. Each image collected had a frame size of 256 by 256 pixels and was scanned at a rate of 25.21 μs/pixel. FLIM data was acquired with an ISS A320 FastFLIM box.

Traditionally, calculating fluorescence lifetimes requires mathematically fitting a series of exponential functions to the intensity decay curve. This approach is computationally difficult and makes assumptions about the raw data. In the phasor approach to FLIM, the Fourier transform of the fluorescence decay curve within each pixel of an image is used to generate coordinates in a polar phasor space without complex fitting routines [18]. Once in this transformed phasor space, the average fluorescence lifetime can be determined (Figure 1). Using the property of linear combinations on this polar plot allow for the quantification of relative amounts of the two species within each pixel. In this study, the relative amounts of free and bound NADH within each pixel of a fluorescence image were quantified using this method. Previous research has suggested that the free and bound forms of NADH reflect the predominant form of metabolism as glycolysis or oxphos, respectively [19]. Therefore, FLIM can be used to assess the metabolic state in the cell by monitoring the relative ratio of the free and bound forms of NADH. Data was processed and analyzed using the fluorescence imaging software, SimFCS available at http://www.lfd.uci.edu/globals/.

**Figure 1.**
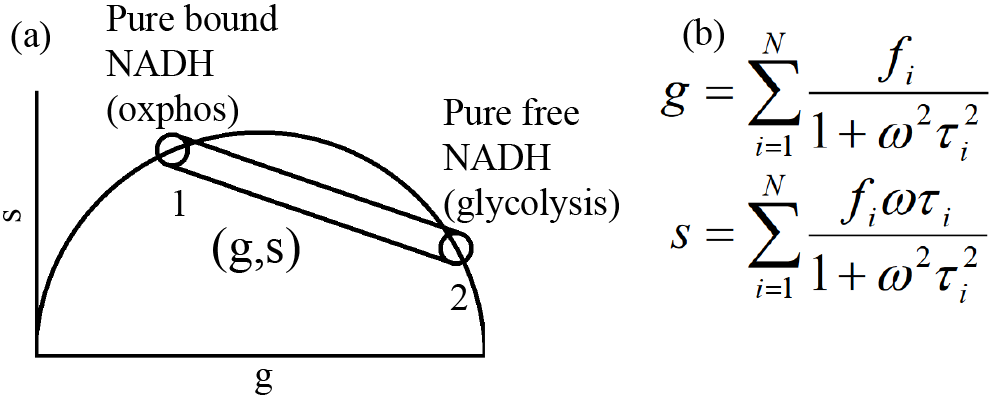
Assessing metabolism using FLIM. (a) The phasor plot allows for visualization of fluorescence lifetimes. Points on the semicircle represent pure lifetimes. The line through them represents possible bound to total NADH ratios. (b) Equations for determining the phasor coordinates for each pixel as a relative fraction of pure lifetime species [18]. The variable *f* represents the relative contribution of a particular species to the phasor coordinates.

### Confocal Laser Scanning Microscopy

In order to visualize the cells and acquire data, experiments were performed on a confocal Zeiss LSM710 with a water immersion 40x/1.2 NA objective. This allowed brightfield and fluorescence images to be acquired on the same system with high spatial resolution. In the laser scanning system, a series of mirrors directs the laser to scan each pixel in a row, moving from left to right, then moves to the next row at the leftmost pixel and repeats this process for all rows. Any fluorescent molecules in the path of the laser would be excited with the proper wavelength. Green fluorescent protein (GFP) excitation was achieved using a one-photon argon ion laser at 488nm. Fluorescence lifetime imaging microscopy data was acquired using two-photon excitation at 740nm in order to collect NADH emission.

### Cell Viability Assay

Cells were plated onto gridded imaging dishes to determine cell survival following cisplatin treatment. Brightfield images of cells on the grid were acquired before cisplatin treatment and after 24 hours after continuous exposure to cisplatin. Cells were counted if they were morphologically normal (flat and noncircular). The cell numbers in the grids were counted both before and after damage and the ratio of surviving cells to initial cells was calculated.

## Results

### p53 promotes oxidative phosphorylation

Fluorescence intensity and lifetime data of NADH in H1299 cells were acquired. Up to 40 frames were averaged in order to increase the signal to noise ratio of NADH fluorescence and to gather a sufficient number of photons to generate the fluorescence lifetime decay curves. Brightfield images of cells were immediately acquired after the acquisition of fluorescence intensity and lifetime data and were used for evaluating cell morphology. The cells that were selected for data collection were chosen to best match normal H1299 cell morphology and had comparable levels of fluorescence intensity expression. For p53-GFP cells, selected cells also had localized GFP in the nucleus and not distributed in the cytoplasm, as p53 is primarily found in the nucleus. The fluorescence intensity images were then analyzed through masking sub-cellular compartments (nucleus and cytoplasm) and determining the ratio of substrate-bound NADH to total NADH using SimFCS software. The ratios were then averaged for each time point. Standard deviation was also calculated and graphed as error bars and p-values were obtained through the Student’s two-tailed homoscedastic t-test.

The p53-GFP H1299 cells had a significantly higher bound to total NADH ratio than the EGFP H1299 control cells (Figure 2). This difference was observed in both the nucleus and cytoplasm. The higher fraction of bound NADH suggests that p53-GFP cells utilize more oxphos for metabolism compared to EGFP cells, demonstrating that p53 promotes oxphos in H1299 cells.

**Figure 2.**
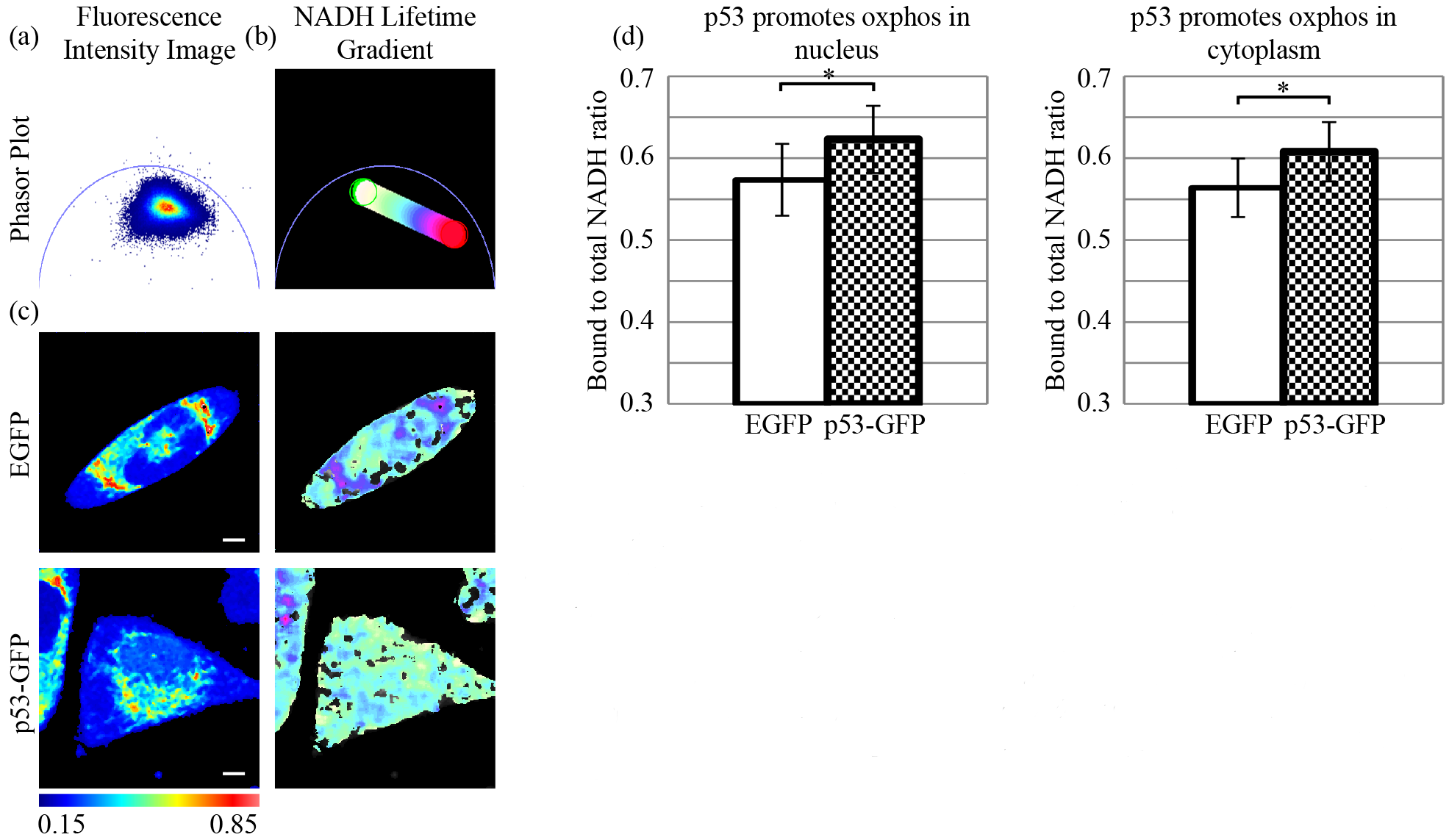
p53 promotes oxidative phosphorylation. (a) The fluorescence lifetime histogram on the phasor plot shows the distribution of lifetimes of autofluorescent NADH. (b) A color spectrum is used to pseudo-color images based on fluorescence lifetime indicating the bound to total NADH ratio. (c) Fluorescence intensity images (left panels) and pseudo-colored FLIM images (right panels) of H1299 lung cancer cells transfected with EGFP (top row) or p53-GFP (bottom row). Red colors indicate glycolytic regions whereas white colors indicate regions of oxphos. Scale bar = 10μm (d) Bar graphs quantifying the bound to total ratio of NADH in the nucleus (left) and cytoplasm (right). Cells expressing p53 exhibit more oxphos.* p<0.05, N>7.

### p53 stabilizes metabolism in response to cisplatin-induced DNA damage

To investigate how metabolism responds to ICLs caused by cisplatin, the fluorescence lifetime of NADH in p53-GFP cells and EGFP cells was tracked before cisplatin treatment and after 12 hours of exposure to cisplatin. The fraction of bound NADH in EGFP cells exhibit sharp decreases in both the nucleus and cytoplasm following cisplatin induced damage, while both the nuclear and cytoplasmic bound to total NADH ratios of p53-GFP cells remain relatively constant (Figure 3). The decrease in both the nuclear and cytoplasmic ratios for EGFP cells are highly significant, while the differences in the average ratio for p53-GFP cells are not statistically significant. The sharp falloff in bound to total ratio of NADH observed in the EGFP cells is not observed in p53-GFP cells, demonstrating that p53 protects oxphos metabolism after damage and prevents the cancer cell from further utilizing Warburg metabolism. Because trends in bound to total NADH ratio in the cytoplasm and nucleus are parallel, further experiments only explored the bound to total NADH ratio in the nuclear compartment.

**Figure 3.**
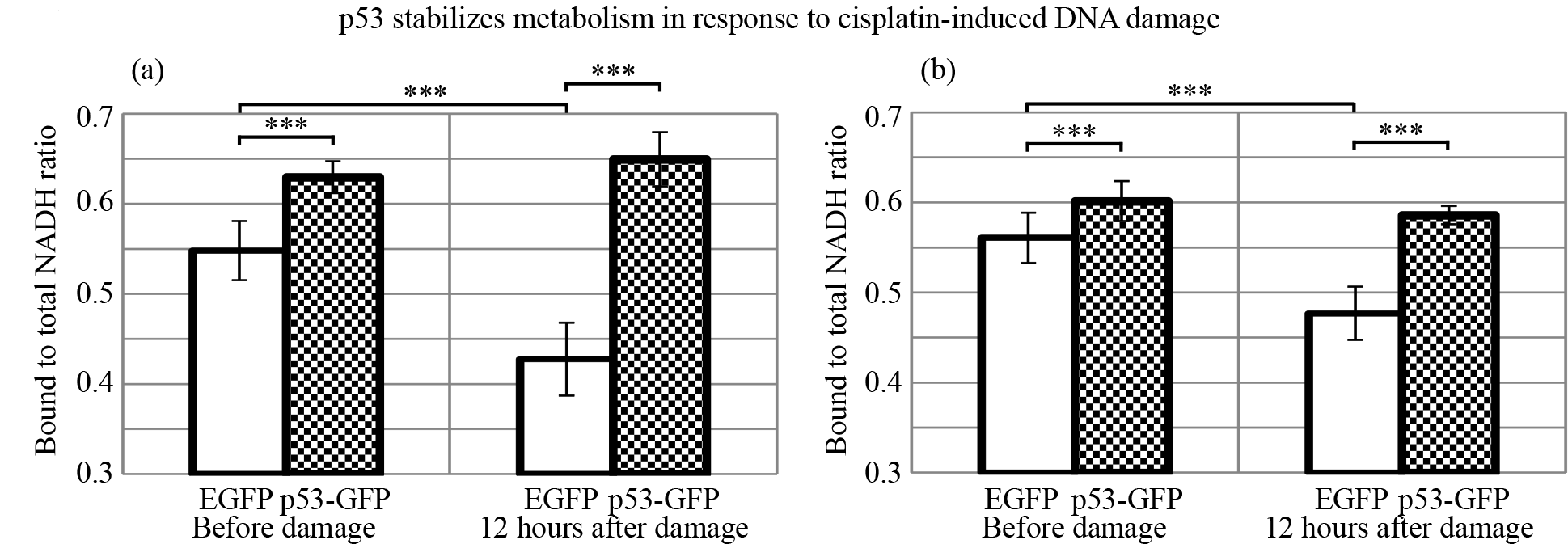
p53 stabilizes metabolism in response to cisplatin-induced DNA damage. (a) Bar graphs of the bound to total NADH ratio before cisplatin treatment and with 12 hours of continuous exposure to 20μM cisplatin in the nucleus and (b) in the cytoplasm. Cells with p53 are able to maintain oxphos usage whereas control cells exhibit a switch toward glycolysis.*** p<0.001, N>7.

### REV3L is necessary for metabolic regulation by p53

Even though the depletion of REV3L in p53-null cancer cells caused an increase in the bound to total NADH ratio in basal conditions, the depletion of REV3L in p53 expressing cells caused a significant decrease of the ratio of bound to total NADH in comparison to p53-GFP cells with functional REV3L (Figure 4). The oxphos pathway used in p53-GFP cells with REV3L depletion was much less utilized than in p53-GFP cells with REV3L prior to damage, indicating that the presence of REV3L is somehow necessary for proper p53 metabolic regulation. The transfection of shRNA or siRNA itself also does not greatly alter the metabolism of H1299 cells as p53-GFP cells with a control transfection still have a significantly greater bound to total NADH ratio than cells that lack functional p53 with a control transfection.

**Figure 4.**
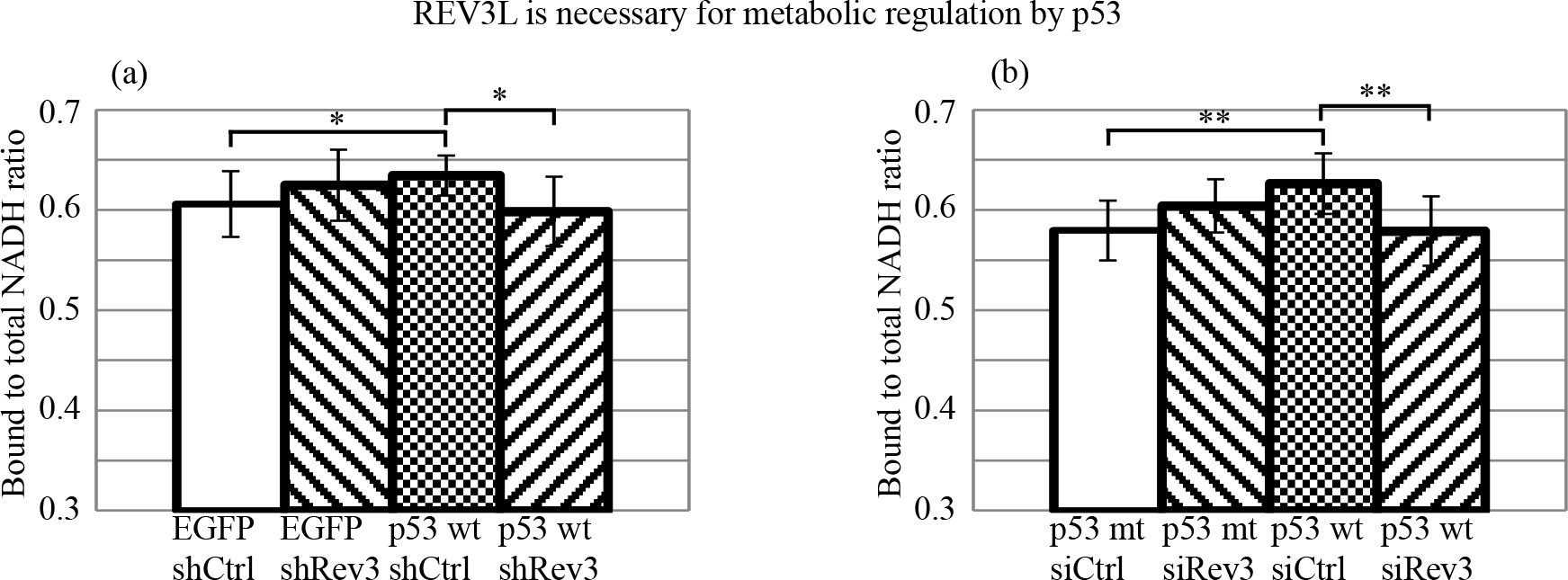
REV3L is necessary for metabolic regulation by p53. (a) Bar graphs of the bound to total NADH ratio in H1299 cells after manipulation of p53 by transfection and REV3L by short hairpin RNA. Cells expressing p53 with silenced REV3L exhibit a lower fraction of bound NADH relative to those only expressing p53 or only expressing REV3L. (b) Bar graphs of the bound to total NADH ratio after transfection of p53-GFP with the R175H mutation and manipulation of REV3L by small interfering RNA. Mutated p53 exhibits similar effects as EGFP expressing cells in (a) showing that functional p53 is necessary for metabolic regulation. Silencing REV3L with either siREV3L or shREV3L results in reduced capacity for oxphos with functional p53. * p<0.05, ** p<0.001, N>7.

### Silencing REV3L and expressing functional p53 increases oxidative phosphorylation following cisplatin treatment

Bound to total ratios of the conditions p53-null (EGFP or mutant p53-R175H-GFP) with control shRNA or control siRNA respectively (the control groups), p53-null with REV3L depletion, p53-GFP expression with control shRNA or control siRNA, and p53-GFP with REV3L depletion were normalized to the corresponding control group ratio at both the before time point and the 24 hours after damage time point (Figure 5). The relative increase was then calculated for each group by dividing the normalized value at the 24 hour time point by the normalized value at the before damage time point. The increase in bound to total ratio was most drastic for the p53-GFP cells with REV3L depletion in both cases and indicates the significant increase in the prevalence of oxphos as a result of the DNA damage caused by cisplatin.

**Figure 5.**
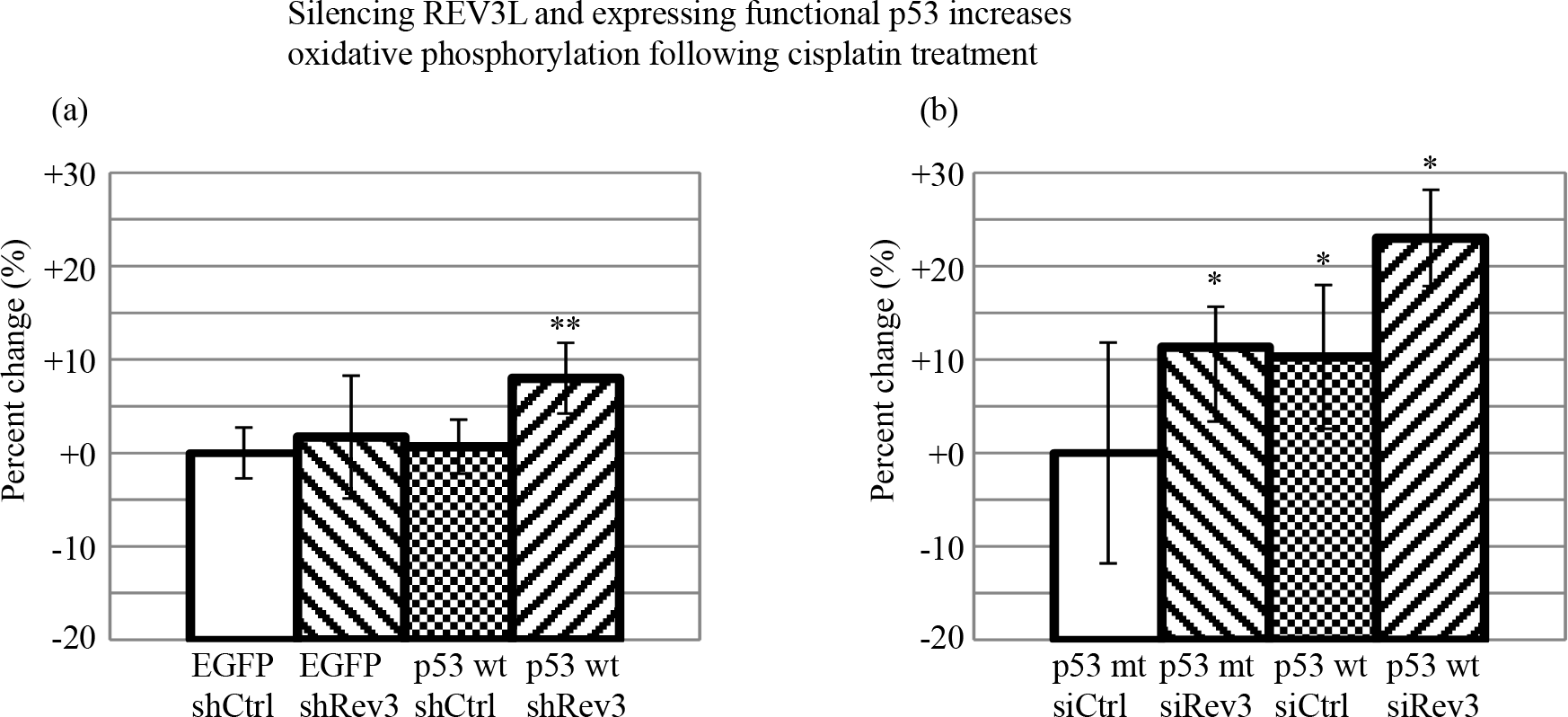
Silencing REV3L and expressing functional p53 increases oxphos following cisplatin treatment. (a) Bar graphs of the percent change in the bound to total NADH ratio after manipulation of p53 by transfection and REV3L by shRNA with 24 hours of 20μM cisplatin treatment. Cells expressing p53 with silenced REV3L exhibit the largest percent change in bound to total NADH ratio representing a metabolic switch to oxphos. (b) Bar graphs of the percent change in the bound to total NADH ratio after transfection of p53-GFP with the R175H mutation and manipulation of REV3L by small interfering RNA. Cells expressing functional p53 with silenced REV3L exhibit the largest percent change in bound to total NADH ratio representing the metabolic switch to oxphos and away from cancer cell Warburg metabolism. * p<0.05, ** p<0.01, N>7.

### Cancer cell proliferation following cisplatin treatment is limited by silencing REV3L and expressing functional p53

We aimed to observe whether or not more cells died following damage for p53 expressing and REV3L depleted cells, corresponding to the idea that for cancer cells non-Warburg metabolism indicates less viability. Cells were plated onto gridded imaging dishes and transfected. 24 hours following transfection, grids were selected for acquiring images, then cisplatin was introduced. Images were then once again taken 24 hours following cisplatin induced damage. The cells that were counted before and after damage were morphologically healthy to ensure that dead cells would not be counted and affect calculations. The cell survival ratio was then determined by comparing how many cells were present in that particular grid after damage and before damage. Cells with silenced REV3L and functional p53 had the least amount of cell survival, followed by REV3L expressing and p53 expressing cells. Cells without any p53, both REV3L depleted and expressing, had the largest amount of cell survival (Figure 6).

**Figure 6.**
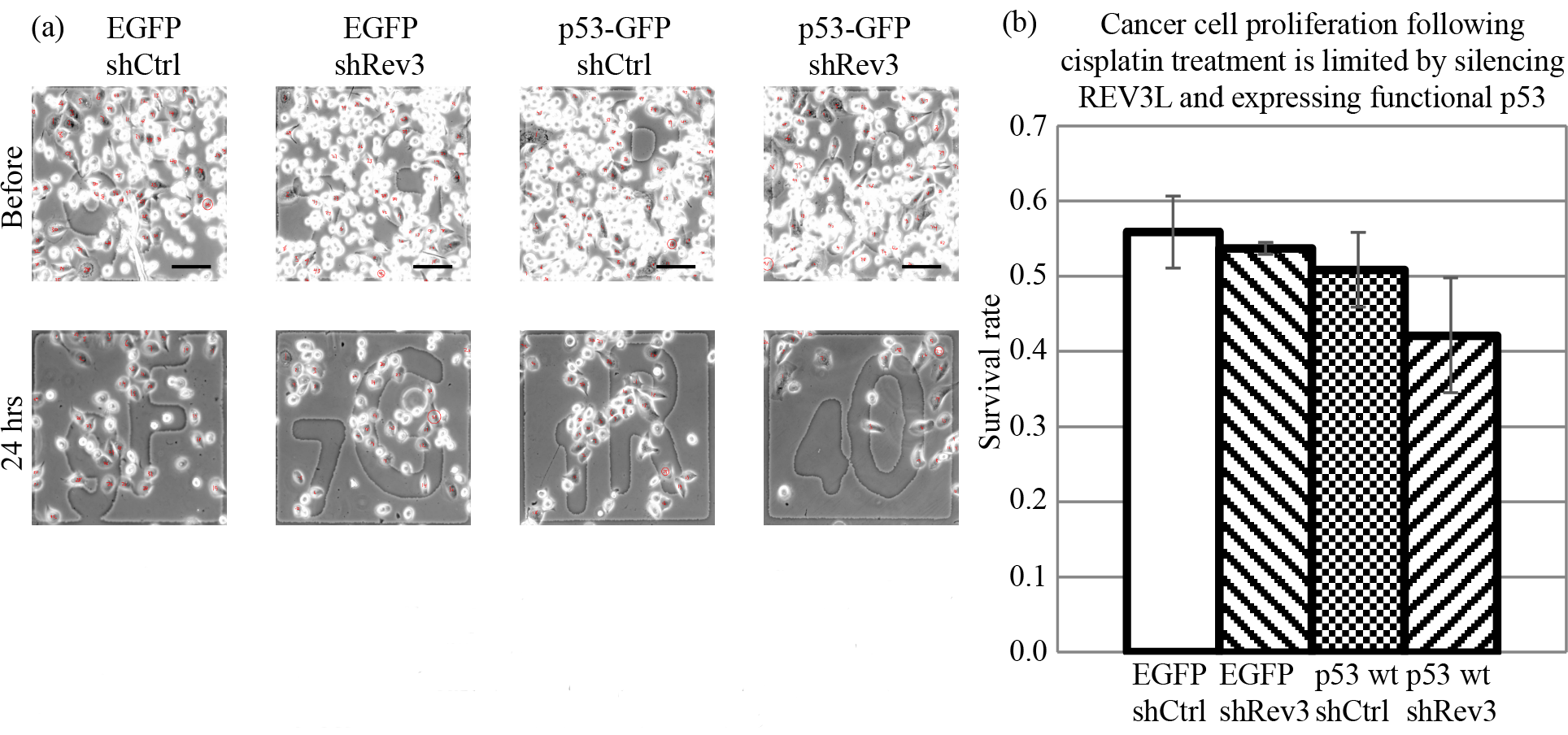
Cancer cell proliferation following cisplatin treatment is limited by silencing REV3L and expressing functional p53. (a) Brightfield images of cells after manipulation of p53 by transfection and REV3L by shRNA with 24 hours of 20μM cisplatin treatment. Scale bar = 100 μm (b) Bar graphs of the cell survival rate of H1299 cells. Cells expressing functional p53 with silenced REV3L exhibit the lowest survival rate after 24 hours of 20μM cisplatin treatment.

## Discussion

The FLIM results suggest that the greatest suppression of the cancerous metabolic phenotype occurs when both p53 function is restored and REV3L is depleted in the presence of cisplatin induced DNA lesions. By depleting REV3L, TLS and the subsequent repair of cisplatin-induced lesions is greatly reduced. Furthermore, DNA damage signaling can then persist. Restoring p53 function in H1299 cells halts the cell cycle and upregulates oxphos, antagonizing Warburg glycolysis. This combination resulted in both the highest bound to total NADH ratio following cisplatin treatment and the greatest percent increase from basal conditions, suggesting that the cancerous metabolic phenotype is severely limited. This presents a viable option for future development of a treatment that can specifically take advantage of the limited Warburg metabolism. The data are corroborated by the low cell survival rate of p53 expressing and REV3L depleted cells after cisplatin damage. As a result of this study, a model for the roles of p53 and REV3L in the cellular metabolic response to cisplatin treatment is proposed (Figure 7).

**Figure 7.**
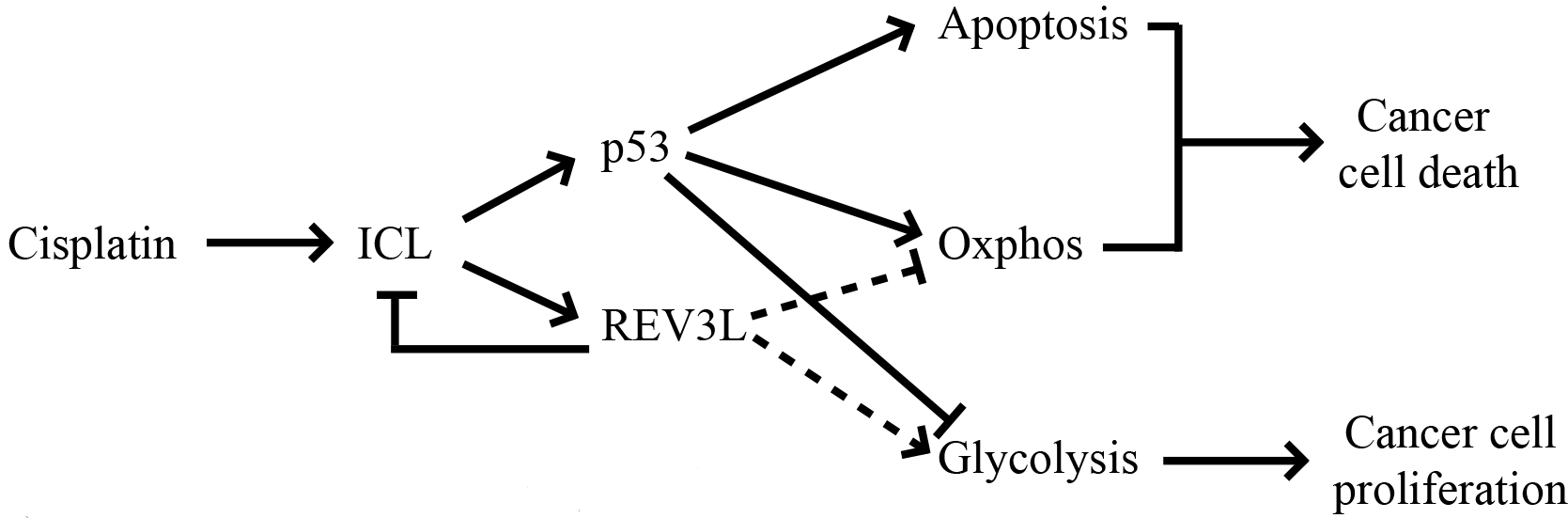
Proposed model for p53 and REV3L activity in modulating cancer cell metabolism following cisplatin damage.

When REV3L is depleted without functional p53, cellular metabolism cannot be regulated by p53. Here, we demonstrate an increase of bound NADH and thus metabolic regulation without p53 activity. The results can be explained by other DNA damage signaling events namely PARP1 activity, which consumes NAD+ and inhibits glycolysis, causing an overall increase of bound NADH [20]. Thus, only depleting REV3L without restoring p53 function can still initiate DNA damage signaling that will ultimate cause suppression of Warburg glycolysis. Developing a molecule to inhibit REV3L is a possible alternative to use in conjunction with cisplatin chemotherapy to prevent the onset of TLS when p53 function is not able to be restored. However, depleting REV3L without restoring p53 function appears less ideal than depleting REV3L with p53 as the survival rate for such cells is very comparable to that of REV3L expressing cells without any p53 function.

The introduction of functional p53 into the H1299 cancer cells expressing REV3L confers the tumor suppressive properties of p53, including its metabolic regulation, which were previously not present. In basal conditions, p53 naturally upregulates oxphos and downregulates glycolysis. Thus, the increase in bound to total ratio seen in the p53-GFP cells when compared to EGFP cells is consistent with previous research [21]. In the presence of damage by cisplatin, DNA damage signaling still occurs, even though TLS through REV3L repairs the lesions, because of ATR signaling pathway among others [22]. This signaling can also cause p53 to increase oxphos and therefore increase the bound to total ratio of NADH. This increase in oxphos in both basal and damaged conditions indicates a suppression of Warburg metabolism, demonstrating that restoring p53 presents a very valid option for co-treatment with chemotherapy in terms of both metabolism and the other tumor suppressive functions that p53 has. However, the process of restoring function of p53 cancers with severe p53 mutations presents an immense challenge itself, but will be very effective in the elimination of cancer cells.

The metabolic regulatory functions of p53 have only been recently discovered within the past decade. Further, the role of REV3L in metabolism is still currently unclear. This study provides some possible insight into the role of REV3L in metabolism and adds another facet to the repertoire of p53 metabolic regulation, namely that in basal conditions REV3L seems to be necessary for optimal p53 regulated oxphos. This is intriguing since it seems to be a novel discovery and links p53 and REV3L in metabolic function.

## Conclusions and Future Work

Overall, the findings of this study suggest that the combination of REV3L depletion, p53 expression, and cisplatin induced DNA damage promotes the most drastic increase of oxphos. This increase of oxphos suppresses the cancerous metabolic phenotype characterized by Warburg glycolysis and presents a very viable option to reduce chemotherapeutic resistance and improve patient outcomes. Here, we demonstrate that p53 expression and REV3L depletion results in the lowest cell survival rate in H1299 cells treated with cisplatin.

In order to improve the current experimental design, fluorescence lifetime data could be tracked over a longer time to observe how metabolism changes further and any long-term effects on cellular morphology and survival. This will more closely model patient responses, as rarely does cisplatin reach peak effectiveness within the first 24 hours in vivo. An increased number of cells for analysis could also be collected in order to obtain a more representative understanding of metabolism. Alternative techniques could be used to support the findings. One such technique is Agilent Seahorse XF technology, which measures O_2_ consumption and H^+^ production to determine oxphos and glycolysis, respectively. Western blots, which determine protein expression through gel electrophoresis and antibody staining, can be used to verify successful depletion of REV3L for siREV3L and shREV3L or no change for siControl and shControl. Cell viability assays through the staining of dead cells can also be done to support the cell survival ratios, as cell death was determined solely by morphology in this study.

In order to develop treatments for cisplatin resistant cancers, a method of restoring the tumor suppressor function of p53 in cancer cells that lack such function should be developed. Finding a way to inhibit REV3L function in vivo should also be a priority for future research, as depletion of REV3L in cancer cells without any functional p53 increased oxphos, albeit to a lesser degree than the combination therapy mentioned above. One of the most interesting additions to the knowledge base of p53 metabolic regulation is the finding that REV3L is necessary for such regulation, a novel discovery. This shows a previously unknown connection between p53 and REV3L in metabolism which requires further research in this fight against chemotherapeutic resistance.

